# A Genotype-to-Phenotype Modeling Framework to Predict Human Pathogenicity of Novel Coronaviruses

**DOI:** 10.1101/2021.09.18.460926

**Authors:** Phillip Davis, Joseph A. Russell

## Abstract

Leveraging prior viral genome sequencing data to make predictions on whether an unknown, emergent virus harbors a ‘phenotype-of-concern’ has been a long-sought goal of genomic epidemiology. A predictive phenotype model built from nucleotide-level information alone has previously been considered un-tenable with respect to RNA viruses due to the ultra-high intra-sequence variance of their genomes, even within closely related clades. Building from our prior work developing a degenerate k-mer method to accommodate this high intra-sequence variation of RNA virus genomes for modeling frameworks, and leveraging a taxonomic ‘group-shuffle-split’ paradigm on complete coronavirus assemblies from prior to October 2018, we trained multiple regularized logistic regression classifiers at the nucleotide k-mer level capable of accurately predicting withheld SARS-CoV-2 genome sequences as human pathogens and accurately predicting withheld Swine Acute Diarrhea Syndrome coronavirus (SADS-CoV) genome sequences as non-human pathogens. LASSO feature selection identified several degenerate nucleotide predictor motifs with high model coefficients for the human pathogen class that were present across widely disparate classes of coronaviruses. However, these motifs differed in which genes they were present in, what specific codons were used to encode them, and what the translated amino acid motif was. This emphasizes the importance of a phenetic view of emerging pathogenic RNA viruses, as opposed to the canonical phylogenetic interpretations most-commonly used to track and manage viral zoonoses. Applying our model to more recent *Orthocoronavirinae* genomes deposited since October 2018 yields a novel contextual view of pathogen-potential across bat-related, canine-related, porcine-related, and rodent-related coronaviruses and critical adaptations which may have contributed to the emergence of the pandemic SARS-CoV-2 virus. Finally, we discuss the utility of these predictive models (and their associated predictor motifs) to novel biosurveillance protocols that substantially increase the ‘pound-for-pound’ information content of field-collected sequencing data and make a strong argument for the necessity of routine collection and sequencing of zoonotic viruses.

## Introduction

To date, the applicability of genomic sequencing data to zoonotic viral outbreaks and pandemics has primarily served in *post*-outbreak genomic epidemiology roles. When a novel viral pathogen emerges, genome sequence data is compared against prior data from other close relatives. From these analyses, public health risk and resourcing (*1*,*2*), transmission chains (*3*), and other response-related information (*4*) is inferred. Several studies have begun to address the utility of viral genome sequencing data in a *pre-outbreak*, predictive methodology through development of increasingly complex machine learning techniques that attempt to understand the emergence of particular viral phenotypes (*5*–*12*). However, while these works provide important novel biological characterization methods, their immediate applied utility for biosurveillance is constrained by limited predictive power and the complexity of interpreting their outputs.

The emergence of the SARS-CoV-2 virus, and the ensuing pandemic, has emphasized our continued vulnerability to zoonotic pathogens. Despite several smaller scale outbreaks of dangerous Betacoronaviruses (namely SARS and MERS), our preparedness and ability to forecast these emergent pathogens have made little advancement. Traditionally, the approach to understanding differences in viral phenotypes has involved problematic experimental evolution, or gain-of-function research through recombinant genetics system (*13*, *14*).

Our previous work developed a feature-agglomeration method adapted to “bag-of-words” style feature extraction in RNA viruses (*15*). We used this method to fit a binary logistic regression model for *Orthocoronavirinae* around a response variable of human pathogen vs non-human pathogen. While this method focused on explanatory modeling by emphasizing numerical stability and training-set accuracy as the model selection criteria, the original feature extraction and model fitting implementation limited its predictive power and resulted in overfitting to the training data. This dilemma of model extrapolation is an old problem in statistical analysis and machine learning (*16*, *17*) and is still salient in biological data science applications. This has led to assertions that the goal of prediction for threat of viral emergence, directly from sequence data, is infeasible based on currently available data and biological knowledge (*18*).

We provide a solution to these problems specifically in the case of *Orthocoronavirinae*, while also demonstrating techniques that could be applied across the viral kingdom. We have developed a protocol for feature extraction and cross-validation that is specific to the viral genomics domain to produce actionable and *predictive* genotype-to-phenotype information for global health and pandemic preparedness experts, directly from genomic data.

## Methods

### Data Labeling and Grouping

We adopted the same data labeling assumptions regarding human-pathogen class membership that were stipulated in our previous work (*15*). To reiterate, bat coronaviruses are assumed to not be human coronaviruses. Civet SARS and camel MERS isolates are labeled as human coronaviruses (reflecting their suspected roles as facilitators of spillover), along with the rest of the known human coronaviruses. All other species of coronavirus are labeled as non-human pathogens.

In the application of group labels for stratified resampling and cross-validation, we created a composite label that combined the species level taxid assigned for each virus sequence with its class label with regards to human-pathogen status. This approach attempts to capture the nuance in certain clades of coronaviruses, such as Betacoronavirus 1, where certain members of the species (e.g., PHEV and Bovine COV) appear to have well defined barriers with regards to their capabilities as human-pathogens but share a species designation with a known human-pathogen coronavirus like OC43 (*19*, *20*). This method results in 63 group labels applied across the training set.

### Feature Extraction

We previously developed a feature extraction method (*15*), *Vorpal*, to reconcile the k-mer-based sequence representations with the inter-example variance in RNA virus genomes. This method worked by counting k-mers across the input sequences, removing k-mers that appear below a frequency quantile threshold, and performing hierarchical clustering on the remaining k-mers. Using hamming distance as the metric and producing flat clusters from the resulting k-mer tree at different branch lengths, we can produce de-facto k-mer alignments that can be re-encoded using International Union of Pure and Applied Chemistry (IUPAC) nucleic acid characters. This functions as a dimensionality-reduction technique that represents the higher dimensional k-mer space into a smaller vocabulary of degenerate motifs that retains information about observed variance in the training data. We can construct feature spaces using this technique that are influenced by three parameters: k size, k-mer frequency cutoff to proceed to clustering, and degeneracy cutoff for flat clustering of the k-mer tree. A simple example to illustrate this concept is depicted in ***Figure 1***.

**Figure 1.**
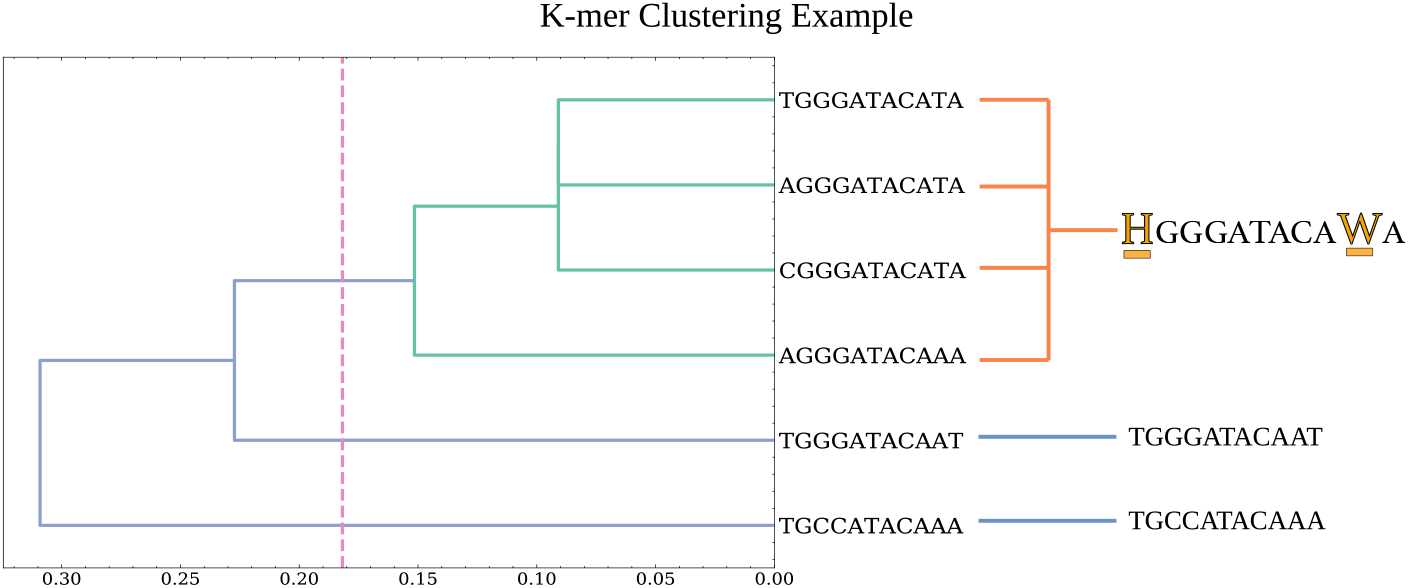
Example clustering of hypothetical 11mers with a 2.0 degeneracy cutoff parameter. Dashed line indicates maximum distance for flat clustering. This distance cutoff is calculated by dividing degeneracy allowance by k-length. In this example 2.0/11 ≈ .182. The four k-mers of the top branches are collapsed into a single ‘degenerate’ k-mer by substituting ‘H’ for the variable T, A, and C bases in the first position and ‘W’ for the variable T and A bases in the second-to-last position.

### Cross-Validation and Resampling

Expanding on this feature extraction technique, we employed several methods to transition this approach from an explanatory paradigm to a *predictive* one. To accomplish this, we utilized two key strategies to reduce possible sources of model variance. First, we used a cross-validation technique to guide model selection that leverages the intrinsic modal organization of genomics data imparted by phylogenetic relationships. This characterizes the problem of predictive phenotype modeling as one where generalization of the model would mean maintaining accuracy to a novel mode of the sample distribution, or in other words, a new species or clade of the viral family. Therefore, we leverage taxonomic organization of the training data to implement a group-shuffle-split (GSS) cross-validation approach (*21*). This simulates the problem of having several species of each class represented in the training set and allows a search over model parameters that maximize the ability to generalize to a withheld species in the validation set. In ***Figure 2***, a visualization of this modality in the sample space is demonstrated through a two-dimensional t-distributed Stocastic Neighbor Embedding (tSNE) using the features for the selected model discussed in the Results.

**Figure 2.**
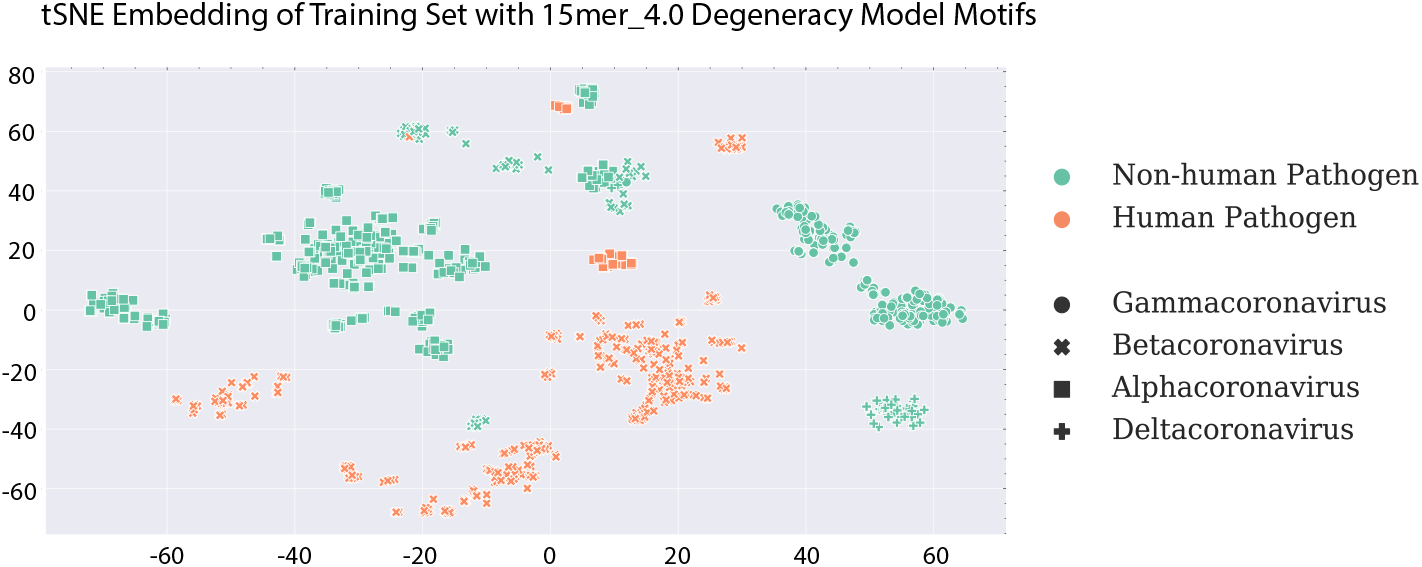
tSNE embedding with features used in the 15mer 4.0 degeneracy-cutoff model examined in results. This visualizes the modality of virus sequences in the sample space.

The second key factor in this predictive modeling approach is the implementation of a stratified resampling technique. Since we chose to use a high-bias model such as logistic regression, what remained was the management of other possible sources of model variance. One substantial source of variance is the skewed representation of complete *Orthocoronavirinae* genomes from clades with clinical and/or other human-related interest. We combat this source of variance by a stratified resampling method (*22*). This resampling method is used at training time to uniformly resample instances from the training set based on the same taxonomic organization utilized in the GSS cross-validation strategy. Additionally, since the *Vorpal* feature extraction methodology is sensitive to this representation bias as a result of the quantile cutoff for k-mer clustering, we use this same resampling technique in the generation of the clustered k-mer motifs. Leveraging this taxonomically-guided resampling at all steps in the process where model variance could potentially be introduced as a side effect of sampling biases allows for effective model training routines to find a closer approximation of the “true” function relating the predictor variables with the response variable.

### Training and Test Set Data

All viral genome sequences for feature extraction and model training were derived from RVDB14, published October 1^st^, 2018 (*23*). Of course, given the publication date cut-off, SARS-CoV-2 records were not present in this data. Additionally, Swine Acute Diarrhea Virus (SADS) sequences were removed from the training data, while bat-HKU2 sequences were left in and labeled non-human pathogens consistent with the rest of the labeling criteria.

In the generation of the test set, SADS and SARS-CoV-2 sequences were downloaded from NCBI Virus (*24*). We subsampled 10 sequences representing each W.H.O. variant-of-concern (VOC) from these downloaded sequences. The test set was completed by adding the RefSeq SARS-CoV-2 reference sequence as well as WA1, to provide representative diversity of sequences across the duration of the COVID-19 pandemic. A total of 42 complete SARS-CoV-2 genomes comprised the full test set of ‘positive’ examples (i.e., human pathogen class label). A total of 34 complete SADS genomes comprised the full test set of ‘negative’ examples (i.e., non-human-pathogen class label). The designation of SADS as a true negative was supported by the apparent zoonotic barrier between humans and porcine coronaviruses in general, as well as reporting of SADS outbreaks in pig farms in China resulting in no documented human sickness in workers exposed to sick pigs (*25*).

Models were fit in triplicate to estimate variance in model accuracy and test set probability as a result of training set resampling and random initialization of coordinate descent. Parameters for GSS were .10 splits, meaning 10% of groups were separated for validation with each split, with 100 training and validation splits produced for each training session. The training set of 2276 sequences was randomly super-sampled to 4000 instances using the stratified resampling method described above. P-values for coefficients were not estimated, as predictive power to withheld data is the preferred model evaluation criteria in this context.

Model selection was performed by first producing degenerate motifs across combinations of two feature extraction parameters; k-mer size and degeneracy cutoff. Then, each of these feature sets was used to fit models with a grid search cross-validation routine that searched over the L1 regularization parameter *C* using GSS as the cross validator, where *C* is the inverse of the L1 regularization term λ. Quantile cutoff for k-mer clustering was selected for each k-size based on available system memory constraints (2TB) and are stipulated in *Supplementary Table 1*. The complete list of parameters and their values is summarized in ***Table 1***. The best estimator was chosen using mean validation set score, where negative Brier score was the scoring function. Brier score is equivalent to mean squared error when the outcome is a binary probability estimate.

**Table 1.** Parameters for feature extraction and LASSO model hyperparameters. Combinations for k-size and degeneracy cutoff resulted in 15 extracted feature sets. These feature spaces were fit in triplicate with Grid Search over these values for C. This resulted in 45 fitted models for comparison.

| Tested K-mer $k$ size | Tested Degeneracy Cutoffs | Tested L1 Regularization Parameters ($C$) |
| --- | --- | --- |
| 11 | 1.0, 2.0 | .01, 0.1, 1, 10, 100, 1000, 10000 |
| 13 | 1.0, 2.0, 3.0, 4.0 | .01, 0.1, 1, 10, 100, 1000, 10000 |
| 15 | 1.0, 2.0, 3.0, 4.0 | .01, 0.1, 1, 10, 100, 1000, 10000 |

The code for feature extraction and model fitting, training and test data sets, and corresponding metadata can be accessed at https://github.com/mriglobal/vorpal. The repository also contains a persistent version of the down-selected model highlighted in the *Results* (15mer_4.0) and a series of scripts to begin predicting on novel sequences. This software is provided under an MIT license. A complete list of accession numbers contained in the training and test sets can be found at https://github.com/mriglobal/vorpal/tree/master/data in the tab-separated text files containing ‘label’ and ‘group’ assignments for each sequence.

## Results

Following an exhaustive search over feature extraction parameters and the L1 regularization-term hyperparameter, several models were identified that correctly classified the test set at 100% accuracy – specifically, the 15-mer models with 2.0 and 4.0 degeneracy cutoff, and the 17-mer models with 2.0 and 4.0 degeneracy cutoff for k-mer clustering (***Figure 3***).

**Figure 3.**
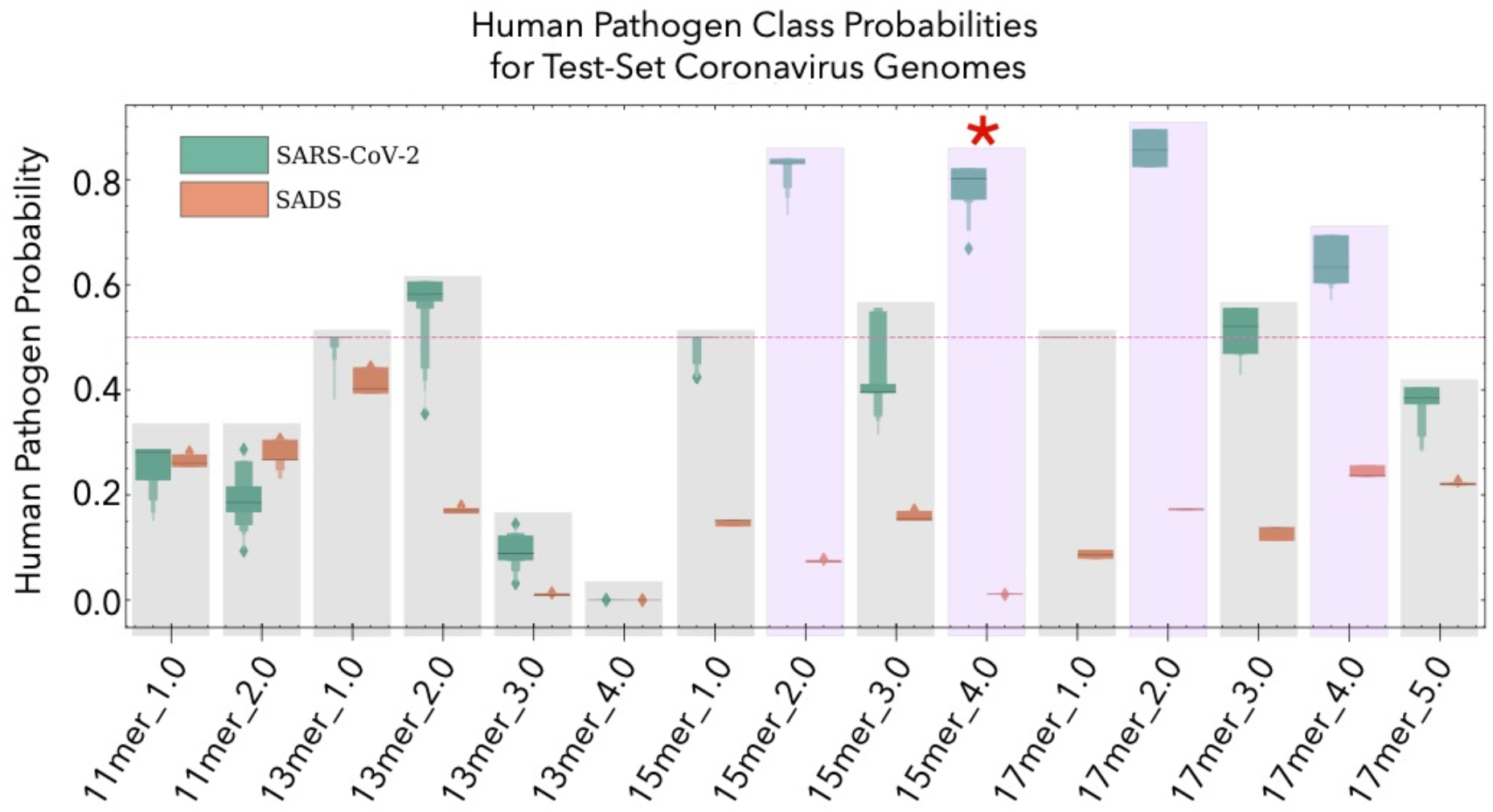
Human pathogen class probabilities for the test set virus genomes across each all model replicates for each combination of feature extraction parameters. Models are titled according to their k-mer length and allowable k-mer degeneracy (e.g., “15-mer_4.0”). The classification threshold of 0.5 is shown as a dashed line. Models that correctly classified all 42 SARS-CoV-2 genome assemblies in the test set as a human pathogen are indicated by light-purple-shaded boxes. Ineffective models are gray-shaded. All models correctly classified all 34 SADS test-set genome assemblies as a non-human pathogen. Red asterisk identifies the feature extraction parameters from which the selected model described in the Results was drawn.

Parameter search over L1 regularization terms was similar to our previous effort. Uniformly, models were selected by the Brier score criterion (*26*) for the strongest regularization term evaluated, which was .01. In *Supplementary Figure 1*, mean cross-validation score is shown to reach an inflection point at this value across all models. We selected one of the 15mer 4.0 model replicates to examine the model predictors in further detail and to provide an example of insights that might be gained from their interpretation about the genomic determinants of human pathogenicity in coronaviruses. Predictor motifs and their corresponding coefficients are provided in ***Table 2***. The coefficients in logistic regression can be interpreted as the linear effect of each unit of the predictor variable on the log-odds of the response variable.

**Table 2.**
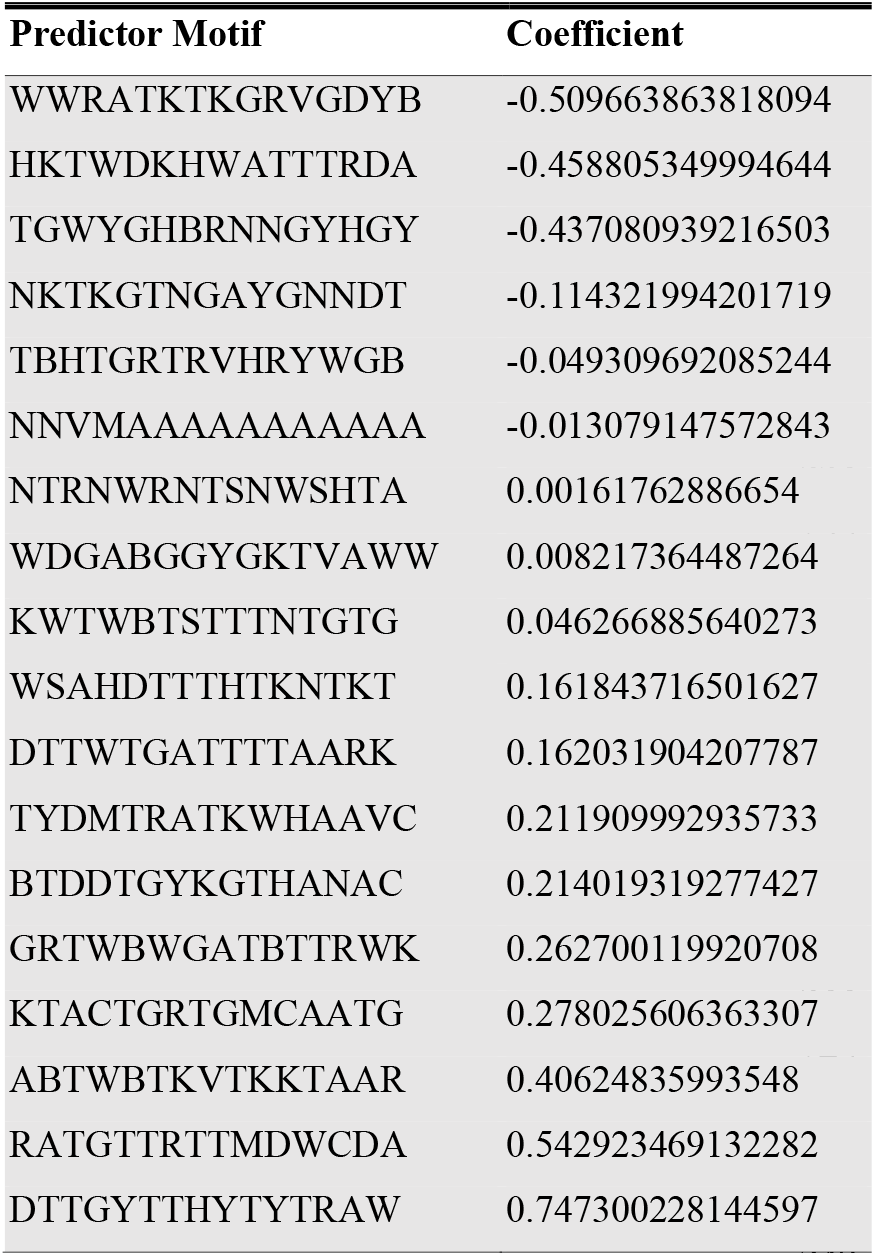
Predictor motifs with nonzero coefficients after LASSO feature selection for the selected 15-mer with 4.0 degeneracy model. Positive coefficients correspond to an increase in the probability of human-pathogen class membership.

A comparison between different coronaviruses and their respective utilization of the predictor motifs allows for interpretation of the functional origin. As an example, ***Table 3*** provides a comparative mapping of the model predictor motif, RATGTTRTTMDWCDA, across a variety of coronavirus species, the corresponding codons for that motif in its genomic context, and the amino acids encoded. The first observation is that these motifs appear in association with human pathogenicity across distantly related coronaviruses across several genera, but in varied genomic loci. Secondly, some motifs provide increased class probability mostly through a binary presence/absence (e.g., DTTGYTTHYTYTRAW), while others, such as NTRNWRNTSNWSHTA, act through frequency enrichment, appearing up to 45 times in some human-pathogen HKU1 isolates and as few as 4 times in Sparrow Coronavirus HKU17. The reuse of these motifs in various genomic contexts, while remaining consistently associated with human pathogenicity in these viruses, suggests phenetic similarity in their function, as well as underscores the importance of the alignment-free characterization of the prediction problem in identifying these phenomena.

**Table 3.**
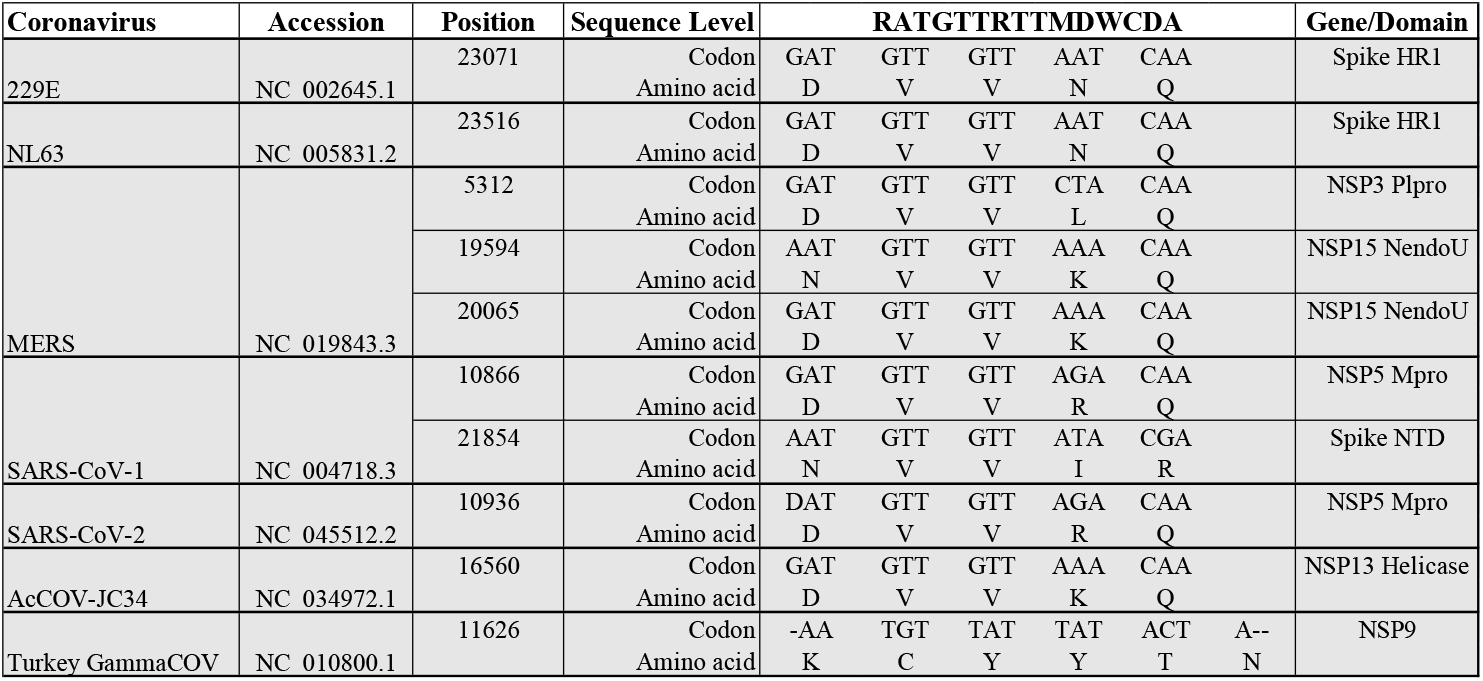
Predictor motif ‘RATGTTRTTMDWCDA’ and the various genomic contexts in which it appears across Alpha-, Beta- and Gammacoronaviruses. The motif always appears in the same reading frame in Alpha- and Betacoronaviruses, while it appears in the +1 position in Turkey Gammacoronavirus (a non-human pathogen).

Interpretation of misclassified instances in the training set, especially for the models that correctly classified the test set sequences, show several interesting patterns. First, proximal phylogenetic ‘near-neighbors’ of known coronaviruses are also proximal in terms of class probability. For instance, WIV16 (*27*), which shares >96% sequence identity to SARS-CoV-1, has a class probability of 0.78 while the civet SARS examples like HC/GZ/32/03 have class probabilities of 0.89 (*Supplementary Data*). This trend continues with late-SARS isolates such as WHU having a predicted class probability of 0.95. Of course, this relationship would be expected for the training data, but this relationship is maintained in the new SARS-CoV-2-related sequences published since the beginning of the pandemic. We used an ensemble of all of the models identified in ***Figure 3***, specifically the 15mer 2.0, 15mer 4.0, 17mer 2.0 and 17mer 4.0 models that successfully generalized to the test set, to predict the class probabilities of these, as well as other novel coronavirus sequences published throughout 2020 and 2021. These results are summarized in ***Figure 4***. Bat coronaviruses with proximity in sequence identity to SARS-CoV-2 (*28*, *29*, *30*), such as BANAL-52, RmYN02, RpYN06 and RaTG13, exhibit human pathogen class probabilities that are proximal to the class probability of SARS-CoV-2. However, while SARS-CoV-2 has the highest mean class probability, it is interesting to note that BANAL-52 has the highest overall sequence identity to SARS-CoV-2 at 96.8%, but is not the closest to SARS-CoV-2 in terms of mean human pathogen class probability.

**Figure 4.**
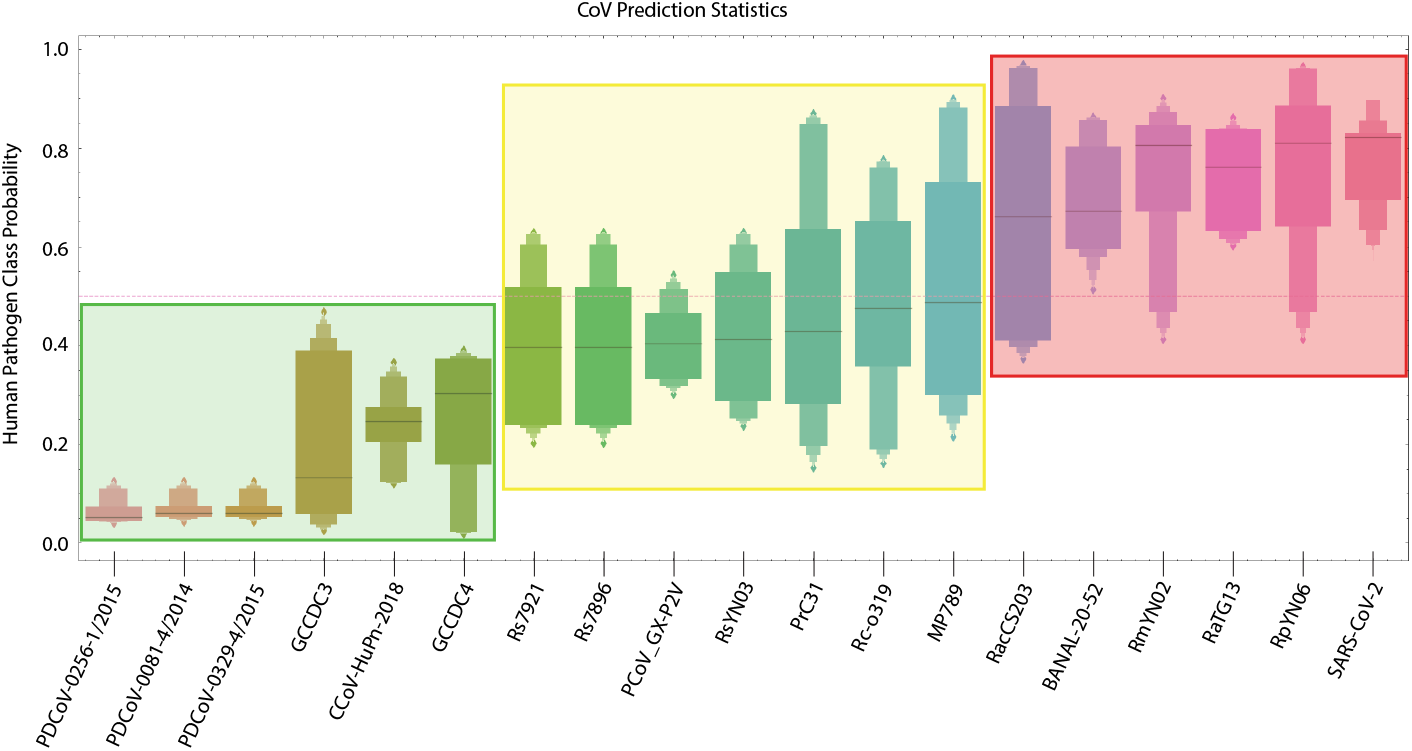
Human pathogen class probability predictions from an ensemble of models identified in Figure 3. The dashed line depicts the classification threshold of 0.5. Red-shaded box indicates viruses with mean human pathogen class probabilities across all models above 0.5. Yellow-shaded box indicates those viruses approaching 0.5 threshold. Green-shaded box indicates viruses well below threshold and unlikely to be a human pathogen according to the plurality of the models.

Another interesting pattern observed in the training set was a group of Bat SARS-like and MERS-like viruses that were routinely classified as human pathogens – specifically, members of Jinning cave group of viruses such as Rs4231 and Rs4874, as well as the MERS-likes NL13845 and NL140422 sampled from a cave in Guangdong (*31*,*32*). These class designations seem to be supported by serological evidence of positivity to SARS-likes reported in the area surrounding the Jinning cave from which these SARS-like viruses were sampled (*31*). Finally, human enteric coronavirus 4408 was classified as a non-human pathogen in 35 of the 45 trained models, including those that were 100% accurate on the test set. Complete tables of misclassified training set accession numbers and class probabilities for each model replicate are available in Supplementary data. The frequency of this misclassification is potentially explained by 4408’s status as a strictly child-associated coronavirus (*33*). Similarly, the novel Canine alphacoronavirus isolated from a child in Malaysia (*34*) in 2018 shares a similar, negative prediction as can been seen in Figure 4. The implications of this nuance in data labeling and the characterization of the problem as a binary classification are examined in the *Discussion*.

## Discussion

Through examination of the model training results, it is possible to see the key determinants of the success of our approach. First, the choice of model – regularized logistic regression – is critical to the success of the models. The 17mer, 3.0 degeneracy models are examples where the models failed to generalize to the test set, but had highest accuracy scores on the training data (i.e., >99%). Controlling this tendency to overfit, especially where certain nuance or ambiguity may exist regarding the virus phenotype that is not captured by the binary response variable, is much more difficult to achieve outside of high bias model families like generalized linear models. Second, the positionally-independent representation of the feature space provided by the *Vorpal* feature extraction methodology allows for identification of genome thematics that emerge as a result of convergent evolution. Finally, the degenerate characteristic of these motif representations introduced by the k-mer clustering clearly contribute to success in extrapolation. This is explained by observing several instances where the models did not successfully generalize to the test set. In many models that were fit with lower degeneracy cutoff parameters, test set probabilities for SARS-CoV-2 were 0.50 because none of the predictor motifs selected during training mapped to SARS-CoV-2 (***Figure 3***). Higher degeneracy feature spaces still identified predictive motifs, and these motifs continued to be present in the test set.

To understand the underlying biological function of the predictor motifs, we examined their genomic context. As an example, RATGTTRTTMDWCDA, shown in ***Table 2***, is located in both SARS-CoV-1 and SARS-CoV-2 at the domain boundary in NSP5, the Main Protease (Mpro), between the catalytic domain and the dimerization domain. The arginine that is coded for in the motif has been demonstrated experimentally in SARS-CoV-1 as critical to dimerization (*35*). This motif appears a second time in SARS-CoV-1, in the same reading frame, but in the N-terminal domain of Spike protein, at a position immediately following an N-linked glycosylation site. We previously reported the association of N-linked glycosylation sites and motifs explanatory for host isolate phenotypes in Influenza A as a result of host specific rare codon selection (*15*). The identification of both N-linked glycosylation sites and protein domain boundaries as being sites of rare codon enrichment provides evidence of a translational efficiency adaptation to facilitate co-translational machinery (*36*, *37*). The identification of translational efficiency adaptations as critical to viral fitness has started to significantly expand in the scientific literature (*38*–*40*).

Properties of the NTRNWRNTSNWSHTA motif that led to its association with human pathogens are not obvious, but examining its patterns of occurrence provides potential hints. As mentioned, this motif is most abundant in HKU1. However, in addition to this frequency, it also occurs concurrently in the genome with another unique feature of HKU1 for which the functional purpose is not understood – this motif tracks each instance of the Acid Tandem Repeats (ATRs) that occur at varying copy number in the hypervariable region of NSP3 in different strains of HKU1 (*41*). This motif also appears to be tracking the abundance of consecutive third-position-thymine codons. The preference of these codons is a well described phenomenon in coronaviruses, but its functional provenance is not well understood and its enrichment specifically in human coronaviruses has not been described (*42*, *43*).

The models also appear to describe a human-pathogen class definition that only includes viruses that can readily transmit between adults. There are now a series of coronaviruses that appear to have the capability to cause clinical illness in children, but the children act as terminal hosts for the virus. This list now includes Canine Alphacoronaviruses observed in Thailand in the early 2000s (*44*) and Malaysia in 2018 (*34*), Murine Hepatitis Virus detected in SRA datasets from children with febrile illness (*45*), Porcine Deltacoronaviruses in children in Haiti in 2014 and 2015 (*46*), as well as human enteric coronavirus 4408 (*33*).

### Nuance to class labeling

There is also a well-documented divide in the symptomology observed in juveniles and adults for SARS-CoV-2 (*47*), that is partially described by lower permissivity of infection not attributable to ACE2 or TMPRSS2 expression levels (*48*). The models, notably, do not contain predictor motifs that pertain to these child-specific coronaviruses as they are routinely classified as non-human. While we are modeling a binary response variable in this work, where ‘human pathogen’ is the positive class, a more accurate description of the class labels we have applied might include a likelihood of observance. There appears to be some stratification, where sustained transmission of the virus in humans is *de facto* included as part of the phenotype definition. Viruses that may be capable of spilling over into humans, but who are, for the virus, terminal hosts, have genotypic features which are not captured in our models.

### A Flexible Framework

This characterization of the problem of genotype-to-phenotype prediction using this feature extraction and modeling approach has several distinct advantages. First, assumptions are minimized beyond the independence assumption about predictor interactions in the logistic regression model, and that the information about the phenotype can be contained in the window size of K. These models have the ability to learn predictors that could be in both non-coding and coding regions, and a result of either nucleic acid or amino acid selection processes. It requires nothing other than the complete genome sequence of the virus, and reduces uncertainty that may arise as a result of limitations in accurate annotation or functional proteome estimation. While this effort represents a specific procedure with respect to this feature extraction technique, the theoretical framework is one that can be generally applied any response variable of interest. The task for supervised learning on biological sequence data is to transform to a feature subspace where the learner is interpolating over the feature space as it pertains to the response variable, and is no longer extrapolating. We believe these methodologies are applicable not just across the RNA virus *genome* domain, but also across multiple feature spaces such as protein and RNA secondary structure. We will explore this in future work.

### Improving Biosurveillance Protocols

The implications of the models support a potential reimagining of biosurveillance efforts and pandemic prevention. The ability to predict pathogenic phenotypes of viruses well ahead of spillover, directly from sequence data, can enable more effective focusing of resource allocation for ecological monitoring and prevention. The results described in this work are, to our knowledge, the first demonstration of this capability. Determination of the biological function of model predictors may yield a more detailed understanding of why certain organisms, such as Camels and Civets, seem to act as keystone species for the spillover of certain viral families like *Orthocoronavirinae*. This could produce a road map to understand the host genomic determinants that condition these viral genomes for emergence from their natural reservoirs.

Leveraging predictive motifs in field-forward ‘sequence-search’ missions can enable genomic epidemiologists to identify problematic viruses more quickly on site. Despite the criticality of genome assembly and phylogenetic analyses during emerging outbreak scenarios, their cumbersome and time-consuming nature limits the utility and feasibility of sequencing operations in field-forward surveillance efforts and prevents investments in such infrastructure and programs. Predictive motifs can be modeled directly in raw voltage disturbance signals from nanopore platforms (*49*). Searching for predictive motifs from raw electrical signal obviates the need for in-field basecalling, enabling more streamlined field-forward sequencing infrastructure. Such infrastructure can alleviate sample bottlenecks at central reference laboratories and establish a more efficient public health response network.

As the COVID-19 pandemic has made abundantly clear, the time is now for investments in these types of next-generation biosurveillance ecosystems. Predictive feature-extraction genome modeling frameworks, such as those described here, are poised to underwrite this emerging paradigm.

## SUPPLEMENTARY INFORMATION

**Supplementary Table 1.** Training set and test set accuracy across all modeled feature parameter combinations (in triplicate). Pink-shaded are those models that correctly classified all 42 SARS-CoV-2 test-set assemblies as a human pathogen and correctly classified all 34 SADS test-set assemblies as non-human-pathogens.

| k_size | degeneracy | quantile_cutoff | training_set_accuracy | test_set_accuracy |
| --- | --- | --- | --- | --- |
| 11 | 1.00 | 0.80 | 0.99 | 0.45 |
| 11 | 1.00 | 0.80 | 0.98 | 0.45 |
| 11 | 1.00 | 0.80 | 0.99 | 0.45 |
| 11 | 2.00 | 0.80 | 0.99 | 0.45 |
| 11 | 2.00 | 0.80 | 0.99 | 0.45 |
| 11 | 2.00 | 0.80 | 0.99 | 0.45 |
| 13 | 1.00 | 0.90 | 0.98 | 0.45 |
| 13 | 1.00 | 0.90 | 0.98 | 0.45 |
| 13 | 1.00 | 0.90 | 0.98 | 0.45 |
| 13 | 2.00 | 0.90 | 0.99 | 0.93 |
| 13 | 2.00 | 0.90 | 0.99 | 0.93 |
| 13 | 2.00 | 0.90 | 0.99 | 0.93 |
| 13 | 3.00 | 0.90 | 0.99 | 0.45 |
| 13 | 3.00 | 0.90 | 0.98 | 0.45 |
| 13 | 3.00 | 0.90 | 0.99 | 0.45 |
| 13 | 4.00 | 0.90 | 1.00 | 0.45 |
| 13 | 4.00 | 0.90 | 1.00 | 0.45 |
| 13 | 4.00 | 0.90 | 1.00 | 0.45 |
| 15 | 1.00 | 0.90 | 0.98 | 0.45 |
| 15 | 1.00 | 0.90 | 0.98 | 0.45 |
| 15 | 1.00 | 0.90 | 0.98 | 0.45 |
| 15 | 2.00 | 0.90 | 0.98 | 1.00 |
| 15 | 2.00 | 0.90 | 0.98 | 1.00 |
| 15 | 2.00 | 0.90 | 0.98 | 1.00 |
| 15 | 3.00 | 0.90 | 0.98 | 0.58 |
| 15 | 3.00 | 0.90 | 0.98 | 0.58 |
| 15 | 3.00 | 0.90 | 0.98 | 0.58 |
| 15 | 4.00 | 0.90 | 0.99 | 1.00 |
| 15 | 4.00 | 0.90 | 0.99 | 1.00 |
| 15 | 4.00 | 0.90 | 0.98 | 1.00 |
| 17 | 1.00 | 0.95 | 0.96 | 0.45 |
| 17 | 1.00 | 0.95 | 0.96 | 0.45 |
| 17 | 1.00 | 0.95 | 0.97 | 0.45 |
| 17 | 2.00 | 0.95 | 0.98 | 1.00 |
| 17 | 2.00 | 0.95 | 0.98 | 1.00 |
| 17 | 2.00 | 0.95 | 0.98 | 1.00 |
| 17 | 3.00 | 0.95 | 0.99 | 1.00 |
| 17 | 3.00 | 0.95 | 0.99 | 0.45 |
| 17 | 3.00 | 0.95 | 0.99 | 0.45 |
| 17 | 4.00 | 0.95 | 0.99 | 1.00 |
| 17 | 4.00 | 0.95 | 0.99 | 1.00 |
| 17 | 4.00 | 0.95 | 0.99 | 1.00 |
| 17 | 5.00 | 0.95 | 0.98 | 0.45 |
| 17 | 5.00 | 0.95 | 0.99 | 0.45 |
| 17 | 5.00 | 0.95 | 0.99 | 0.45 |

**Supplementary Figure 1.**
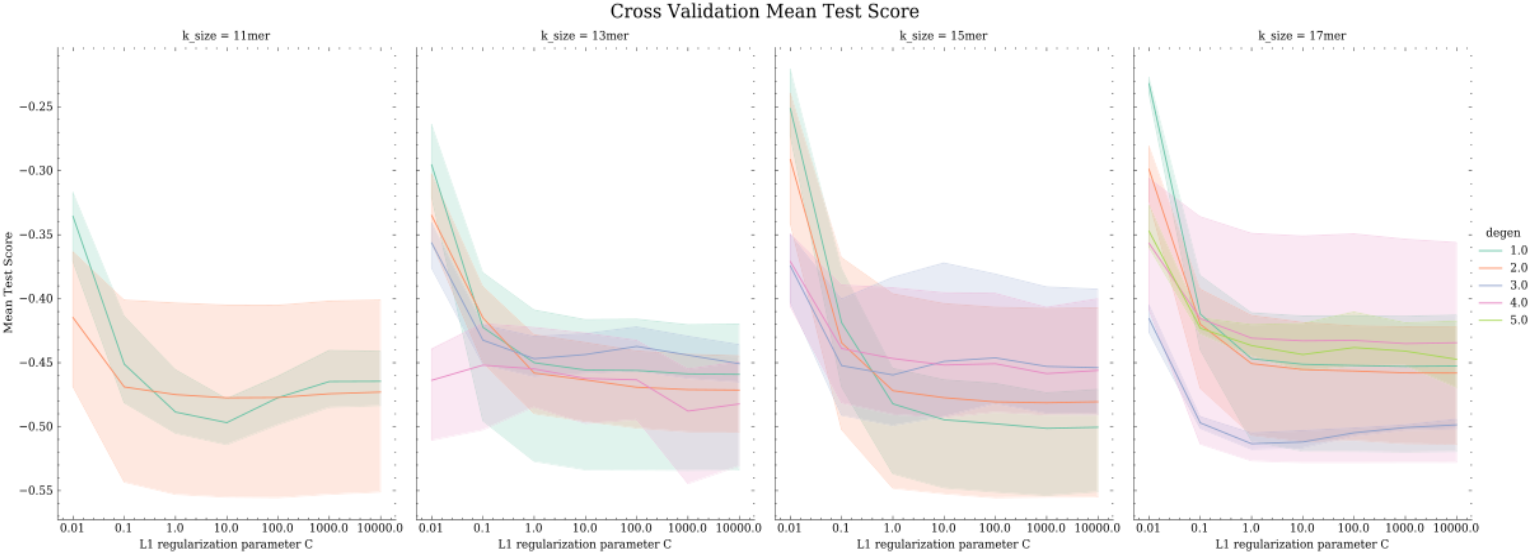
Mean Cross Validation Scores for the validation splits across all k-sizes and degeneracy cutoffs for clustering. Almost every model show dramatic improvements in Brier score as the regularization parameter gets stronger.

